# Ephrin Forward Signaling Controls Interspecies Cell Competition in Pluripotent Stem Cells

**DOI:** 10.1101/2024.06.02.597057

**Authors:** Junichi Tanaka, Yuri Kondo, Masahiro Sakurai, Anri Sawada, Youngmin Hwang, Akihiro Miura, Yuko Shimamura, Dai Shimizu, Yingying Hu, Hemanta Sarmah, Zurab Ninish, James Cai, Jun Wu, Munemasa Mori

## Abstract

In the animal kingdom, evolutionarily conserved mechanisms known as cell competition eliminate unfit cells during development. Interestingly, cell competition also leads to apoptosis of donor cells upon direct contact with host cells from a different species during interspecies chimera formation. The mechanisms underlying how host animal cells recognize and transmit cell death signals to adjacent xenogeneic human cells remain incompletely understood. In this study, we developed an interspecies cell contact reporter system to dissect the mechanisms underlying competitive interactions between mouse and human pluripotent stem cells (PSCs). Through single-cell RNA-seq analyses, we discovered that Ephrin A ligands in mouse cells play a crucial role in signaling cell death to adjacent human cells that express EPHA receptors during interspecies PSC co-culture. We also demonstrated that blocking the Ephrin A-EPHA receptor interaction pharmacologically, and inhibiting Ephrin forward signaling genetically in the mouse cells, enhances the survival of human PSCs and promotes chimera formation both *in vitro* and *in vivo*. Our findings elucidate key mechanisms of interspecies PSC competition during early embryogenesis and open new avenues for generating humanized tissues or organs in animals, potentially revolutionizing regenerative medicine.

## Main

Developmental processes are governed by evolutionarily conserved ligand-receptor interactions^1^. The developmental divergence between distinct species poses challenges for forming chimeric animals during embryogenesis^2–4^. In early embryogenesis, host embryos reject donor human pluripotent stem cells (PSCs) through mechanisms that are not fully understood^4,5^. This rejection may involve various factors such as cell metabolism, interspecific cell competition, and developmental timing differences between humans and host animals^5^. However, the specific signaling pathway that triggers cell death and eliminates human cells in early embryogenesis remains elusive. Overcoming this interspecific barrier is crucial for studying early human development and developing strategies for humanized organ generation via blastocyst complementation in livestock and its future transplantation^6^.

Cell competition is characterized by the death of unfit ’loser’ cells and the survival of fit ’winner’ cells by controlling cell sorting, death, and metabolism, which ensures tissue patterning and integrity^7^. In intraspecies cell competition of drosophila and mouse development, Myc determines the fates of winner and loser cells, while interspecific cell competition depends on the undefined mechanism that activates the p65-MYD88-NFkB-dependent apoptotic pathway in human PSCs^5,8^. Recent studies have demonstrated the potential to complement mesonephros deficiencies in pigs with Six1, Sall1 null renal deficiency by injecting human PSCs overexpressing BCL2 and MYCN using the blastocyst complementation method^9^. Ideally, interspecies cell competition has to be overcome by injection of human cells without overexpression of these tumor-associated genes.

The present study shows the possibility that Ephrin regulates intraspecies cellular competition. The Eph receptor family, as the largest subfamily of receptor tyrosine kinases (RTKs), plays a pivotal role in complex cell communication with its Ephrin ligands, influencing tissue morphogenesis and various pathogeneses^10^. Comprising 14 distinct types, Eph receptors interact with their counterpart ligands, Ephrins, which are subdivided into 5 ephrin-As and 3 ephrin-Bs, each with distinct structural features^10^. This intricate ligand and receptor interaction activates bidirectional signaling in both signal-sending Ephrin-expressing cells (reverse signaling) and signal-receiving Eph-expressing cells (forward signaling), regulating cell migration, adhesion, repulsion, sorting, and apoptosis, potentially crucial for overcoming interspecific barriers^11^.

In this study, we investigated how host cells induce apoptosis in donor cells through cell surface ligand-receptor interaction during interspecies cell competition using an interspecies cell-cell contact-specific reporter system. We identified Ephrin-Eph signaling as a critical regulator in interspecific cell competition, influencing human iPSC cell survival in vitro and in vivo, which is pivotal for humanized chimera formation in mouse development.

### Fine-tuning the interspecies cell-cell contact reporter system

Interspecific cell competition depends on cell-cell contact^5^. However, the mechanism by which mouse cells induce cell death signals in human cells remained unclear. To investigate the potential cellular mechanism, we generated membranous EGFP (mEGFP) expressing human induced pluripotent stem cells (iPSCs) via lentivirus and nuclear tdTomato (ntdTomato) expressing mouse iPSC and performed live-cell imaging in the primed-type PSC maintenance, NBFR medium. Under NBFR medium conditions, human iPSCs maintained pluripotency markers and exhibited prime type gene expression patterns, supported by the previous reports^12^ (Extended Data Fig. 1a). By plating both cells in a sparse cell condition, live imaging revealed that mEGFP^+^ human iPSCs induced cell death upon contacting neighboring ntdTomato^+^ cells (Extended Data Movie 1). This result suggests cell death is induced upon cell-cell contact despite the low cell density, ruling out the possibility of requiring a high cell density-based cell death mechanism via mechanical cell extrusion^13^.

How do mouse iPSCs recognize neighboring human iPSCs and initiate communication to reject them? We hypothesized that mouse iPSCs may send some signaling to the neighboring human iPSCs to induce cell death regardless of cell density.

To test this, we developed an interspecies cell-cell contact-dependent reporter system in which only mouse cells in contact with human cells would express fluorescence. We refined a previously reported synthetic molecule expression construct, composed of SFFV promoter α-GFP- NotchECD-TM-TetR-VP64; anti-GFP-nanobody, Notch extracellular (NotchECD), transmembrane domain (NotchTM), and a targeted molecule (TetR-VP64), which requires tetracycline responsible element-containing promoter (tetO minimal promoter): tetO minimal promoter to turn on the desired gene expression such as BFP^14^. This molecular switch leveraged the Notch ligand-receptor activation mechanism. The anti-GFP nanobodies will bind to the GFP expressed in the neighboring cells and activate the Notch-dependent downstream cascade. The switched-on NotchTM will recruit the gamma-secretase that cleaves the ICD domain of Notch and release TetR-VP64 to express BFP. Any other gene can replace BFP or TetR-VP64 to define the desired gene in this system. To avoid cytotoxic effects of monitoring and to detect early cell-cell contact, BFP for mCherry, a monomeric fluorophore with a relatively quick protein maturation character among the red fluorophores. To test this system, we established surface GFP (sGFP)-expressing human iPSCs via recombinant lentivirus. Then, we examined the efficacy of whether this system would work for interspecific cell communication by using mouse iPSCs expressing a synthetic Notch and sGFP-stably expressing human iPSC co-culture^14^. However, we observed highly leaky mCherry^+^ cells in this classical system even when there was no contact with sGFP-expressing cells (Extended Data Fig. 1b). Despite no contact with GFP, we often kept the significant leakiness of the mCherry signal in this system. We suspected this leakage was due to the unoptimized promoter activity in the Tet-mediated gene regulation system^15^. Thus, we distilled and optimized this system by swapping promoters and succeeded in tightly regulating mCherry expression. We replaced the SFFV promoter with EF1α promoter and the tetO minimal promoter with a Dox-inducible tight promoter (2nd generation of the Tet-Off system)^16^. Based on the high specificity and potential flexibility of this system, we named the construct the “**i**nterspecific cell-cell contact-dependent molecular **<u>switch</u>**(iSwitch).” (Fig. 1a-c).

**Figure 1.**
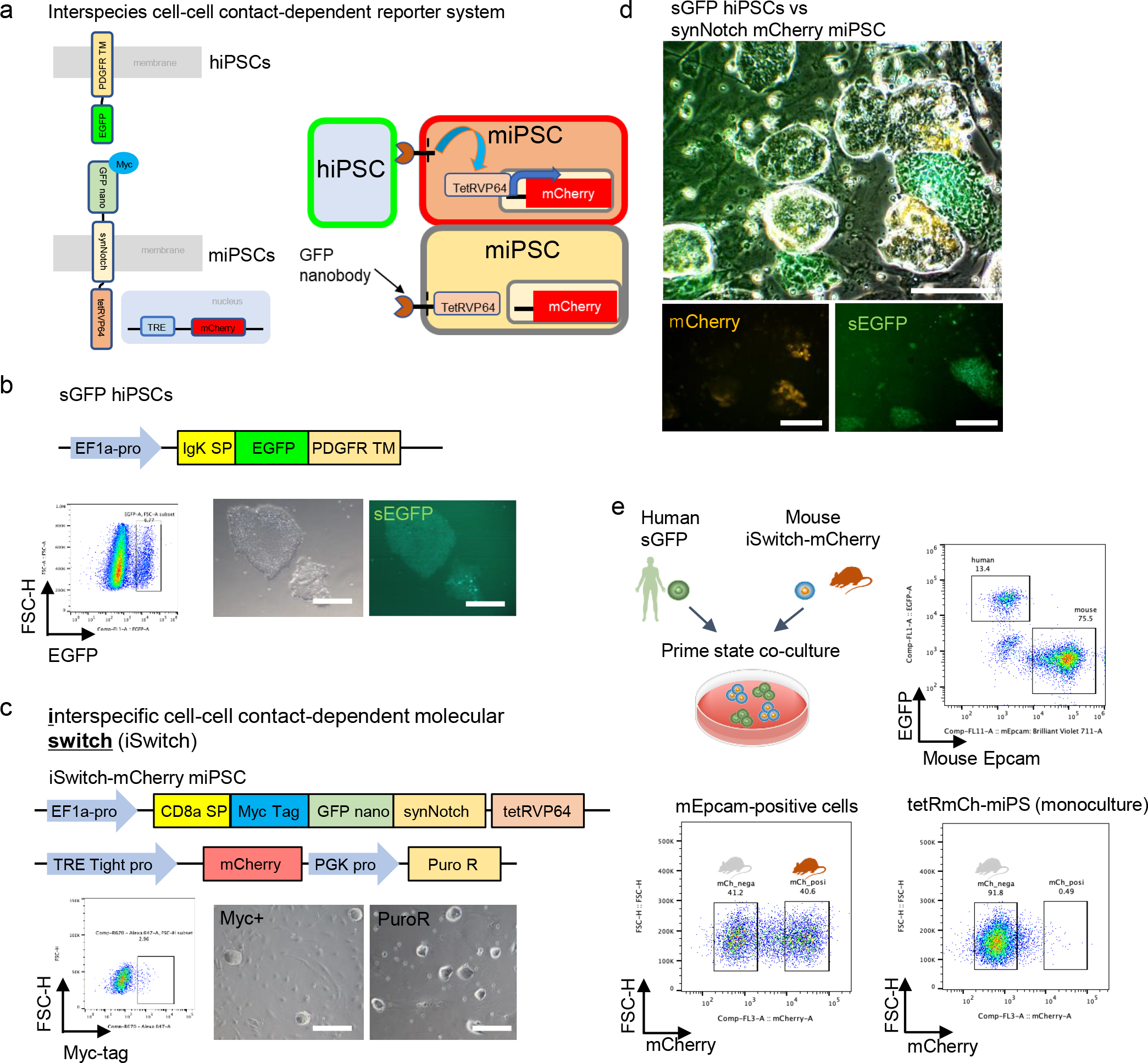
Generation of an interspecies cell-cell contact reporter system. **a**. Schematic of strategy for the cell-cell contact reporter system in human and mouse PSC co-culture. When sGFP-expressing human cells come next, mouse cells express mCherry. **b**. Human cells expressed EGFP on the cell surface using IgK signal peptide and PDGFR transmembrane domain. Lentivirus-infected cells were sorted by flow cytometry. Scale bars, 200Lμm. **c**. For mouse cells, we generated and infected lentivirus for α-GFP-NotchECD-TM-TetR-VP64. Myc-tag Infected mouse iPSCs were sorted by Myc-tag antibody. For the reporter system, Myc-tag-positive mouse iPSCs were infected lentivirus for TRE tight promoter dependent mCherry construct. Infected cells were selected by puromycin. Scale bars, 200Lμm. **d**. Representative microscope images of the iSwitch system of the mouse iPSC contacted with sGFP-positive human PSCs expressed mCherry. Mouse iPSCs that do not come into contact with sGFP-positive human cells do not express mCherry. Scale bars, 200Lμm. **e**. The iSwitch system enables the separation of contact and non-contact mouse PSCs using flow cytometry. Human cells and mouse PSCs were sorted by GFP and anti-mouse Epcam antibodies, respectively. Contact and non-contact mouse PSCs can be separated by mCherry expression. Under mouse monoculture conditions, mCherry-positive cells were few.

We established an iSwitch-stably expressing mouse iPSC line via lentivirus and cultured it in an NBFR medium. The replacements dramatically reduced leakiness and induced the mCherry expression exclusively to the responding neighbors of mouse iPSCs to the sGFP-expressing human iPSCs (Fig. 1d, e). Upon cell-cell contact of iSwitch-expressing mouse iPSC to the sGFP- expressing human iPSCs, the mCherry signal was induced exclusively in mouse iPSCs but not in non-contacting mouse iPSCs, analyzed by fluorescent microscopy (Fig. 1d) and flow cytometry (Fig. 1e). Expression of mCherry in the contacted mouse iPSCs was detected 10 hours after the start of interspecies co-culture (Extended Data Fig. 1c). These results indicate that with this iSwitch system, it is possible to distinguish between mouse iPSCs that initiate interspecies cell-cell contact and those that have not, based on the expression of mCherry.

### Identification of Ephrin A-EphA receptor signaling through ligand and receptor interaction analyses via single-cell RNAseq

To identify the critical signaling pathways regulating the initiation of interspecies cell competition, we performed a co-culture of iSwitch-expressing mouse iPSCs and sGFP-expressing human iPSCs and performed single-cell RNAseq analyses (Fig. 1e). The cell populations were isolated 10 hours post-co-culture initiation through flowcytometry using mouse Epcam antibody (Fig. 1e). The flowcytometry sorted three types of cells: sGFP-positive human iPSCs, mouse Epcam-positive mCherry-negative non-contact mouse iPSCs, and mouse Epcam and mCherry double-positive mouse iPSCs in contact with human iPSCs (Fig. 1e). In the monoculture of mouse iPSCs, mCherry- positive cells were scarcely detected, indicating the appropriate functioning of the interspecies cell-cell contact-dependent reporter system. These three cell populations were subjected to single-cell RNA-sequencing (scRNAseq) analysis.

First, we compared the gene expression profiles of mCherry^+^ mouse iPSCs that contact human iPSCs and mCherry^-^ mouse iPSCs that didn’t (Fig. 2a). The UMAP plot integrating both mouse iPSCs groups showed 14 clusters (Fig. 2b). Cluster 12 was specific to mouse cells in contact, and we identified differentially expressed genes (DEGs) in cluster 12 and performed GO term analysis (Fig. 2c and Extended Data Fig. 2a, b). The results revealed enrichment of apoptosis-related genes such as *Jun* and *Anxa5* in cluster 12 in mouse iPSCs, indicating that even the winning mouse cells in interspecies co-culture can undergo apoptosis. Contrary to expectations, significant gene expression changes in cell groups other than cluster 12 were not apparent on the UMAP between mCherry^+^ responding cells and mCherry^-^ non-responding cells (Fig. 2c). This unexpected result indicates a cell death reaction between mouse-to-human iPSC occurs through intrinsically conserved molecules in mouse iPSC without changing the transcriptional program. Based on this, we performed further computational analyses using CellChat^17^ to identify the ligand-receptor (LR) pairs between sGFP^+^ human iPSCs and mCherry^+^ responding mouse iPSCs, and we used mCherry^-^ mouse iPSCs as a reference (Fig. 2d, e). Although many ligand-receptor interactions were identified, we focused on cell-cell contact-dependent signaling pathways due to the nature of interspecies cellular competition (Fig. 2e and Extended Data Table 1)^5^. Through the CellChat analysis, EPHA signaling emerged as a candidate pathway, particularly the Efna4-EPHA1 and Efna4-EPHA2 signals. Interestingly, the Efna4-EPHA1 signal was anticipated from mouse to human iPSCs, while EFNA4-Epha2 was also predicted from human to mouse iPSCs (Fig. 2f, g). Violin plots showed EPHA1 was explicitly expressed in human cells, whereas Epha2 was predominantly expressed in mouse iPSCs (Fig. 2h). To confirm the certainty and probability of these predicted ligand-receptor interactions from CellChat and narrow down the potential candidate pairs, we also predicted them using scTenifoldXct^18^ (Extended Data Table. 2). Among the candidate ligand-receptor pairs in the CellChat, we checked the common candidate pairs identified both in CellChat and scTenifoldXct. Based on this rigorous selection step, we identified the common ligand-receptor pairs: The Efna4- EPHA1 signal from mouse to human iPSCs and the EFNA4-Epha2 signal from human to mouse iPSCs (Fig. 2i, j). These scRNAseq results suggest a potential role of Ephrin A and EphA receptor signaling in interspecies cellular competition.

**Figure 2.**
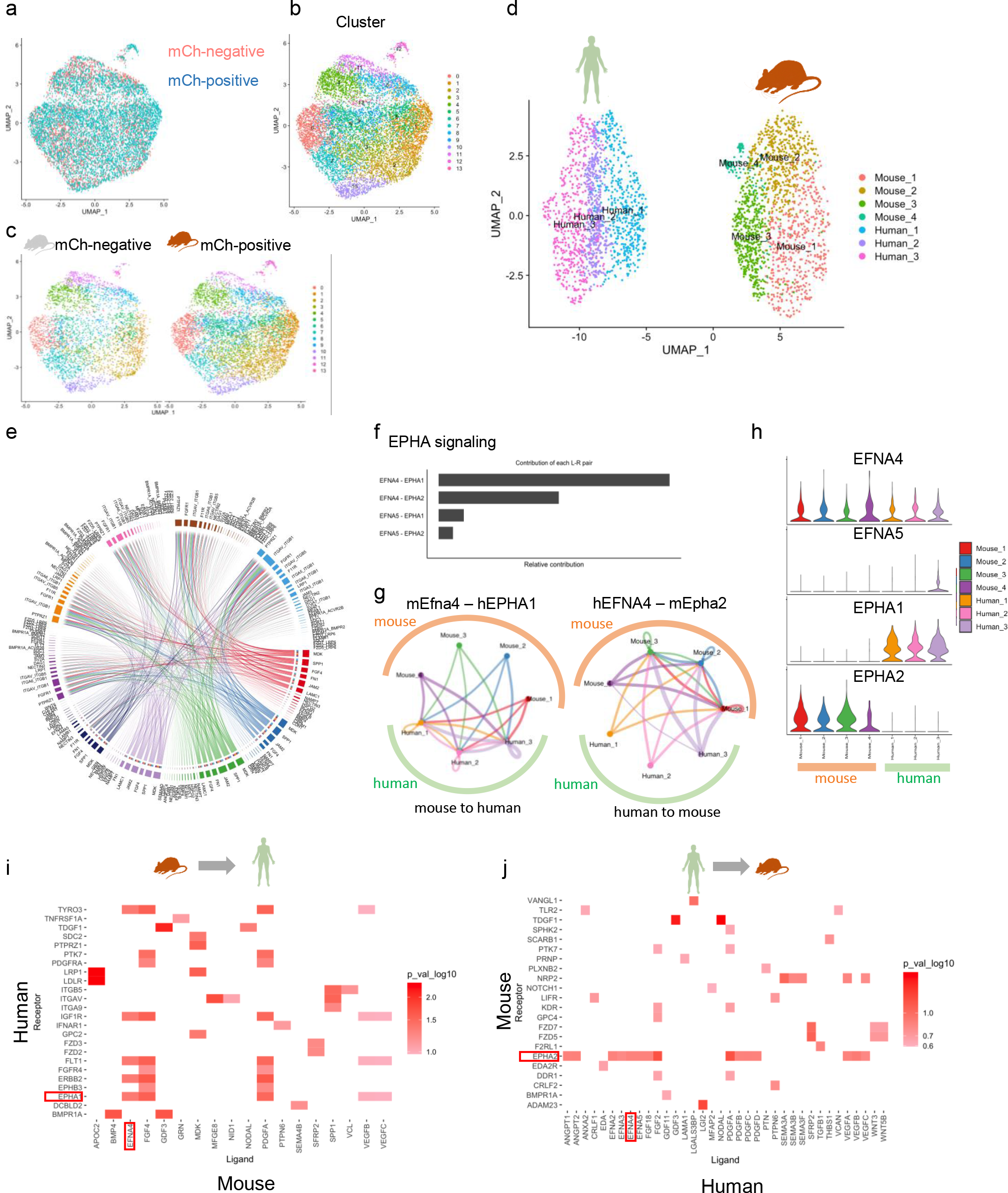
scRNAseq revealed Ephrin A-EphA receptor signaling as a factor in interspecies cellular interactions through ligand and receptor interaction analyses **a**. UMAP showing changes in mCherry-negative non-contacted mouse PSCs and mCherry-positive contacted mouse PSCs. **b**. UMAP shows the cluster of mouse PSCs in co-culture with human PSCs condition. **c**. Separated UMAP showing each cluster was shared by contact and non-contact cells. **d**. UMAP of co-cultured human PSCs and contacted mouse PSCs combined. **e**. Ligand-receptor analysis by CellChat using contacted mouse PSCs and human PSCs. Chord diagrams of mouse PSCs-human PSCs communications. The width of the edge represents the interaction strength. **f**. The relative contribution of each Ligand-Receptor pair to the overall communication network of EPHA signaling. **g**. Circle plots of Ligand-Receptor pairs between different cell clusters. The width of the edge represents the interaction strength. A thicker edge line indicates a stronger signal. **h**. Violin plots showing the expression levels of Ligand-Receptor genes related to EPHA signaling. **i**, **j**. Heatmap shows the relative importance of each Ligand-Receptor pair based on the scTenifoldXct analysis in mouse to human (i) and human to mouse (j).

### Ephrin forward signaling from mouse cells to human cells regulates interspecies cell competition

To validate the results of scRNAseq and ligand-receptor interaction analysis, we inhibited Ephrin-EphA signaling in interspecies co-culture. Since Ephrin ligand-Epha receptors are known to have a high redundancy in their function^19^ and we observed Ephrin and Eph receptors expression in both mouse and human iPSC (Fig. 2h), we first used the pan-antagonist of Eph receptors, UniPR129, that binds in Eph receptor^20^ (Fig. 3a), to inhibit whole Ephrin A-EphA signaling. We analyzed the chimerism of each proportion of human or mouse iPSCs (Fig. 3b, c) and flow cytometry (Fig. 3d). Upon adding UniPR129 to the co-culture of mGFP-expressing human iPSCs and ntdTomato- expressing mouse iPSCs, no difference in cell proportion was observed between the inhibited and control groups after two days of co-culture (Fig. 3b, c). However, from the fourth day of culture onwards, the proportion of human iPSCs decreased in the control group, becoming the loser. In contrast, the proportion of human iPSCs significantly increased in the inhibited group, co- proliferating with mouse iPSCs (Fig. 3b-d and Extended Data Fig. 3a, b). Interestingly, in interspecies co-culture, different species’ colonies typically expand independently, without one species infiltrating the other’s colony (Fig. 3b). However, in the inhibited group, mixed merged colonies of human and mouse cells were notably observed (Fig. 3c). Additionally, adding UniPR129 to monocultures of human or mouse iPSCs showed no significant impact on their proliferation (Extended Data Fig. 3c). These results suggest that Ephrin-Eph signaling inhibition is critical for blocking the cell death of human iPSC in interspecies cellular competition. However, whether this phenotype is initiated from the mouse and human sides or involves forward or reverse signaling remains unclear.

**Figure 3.**
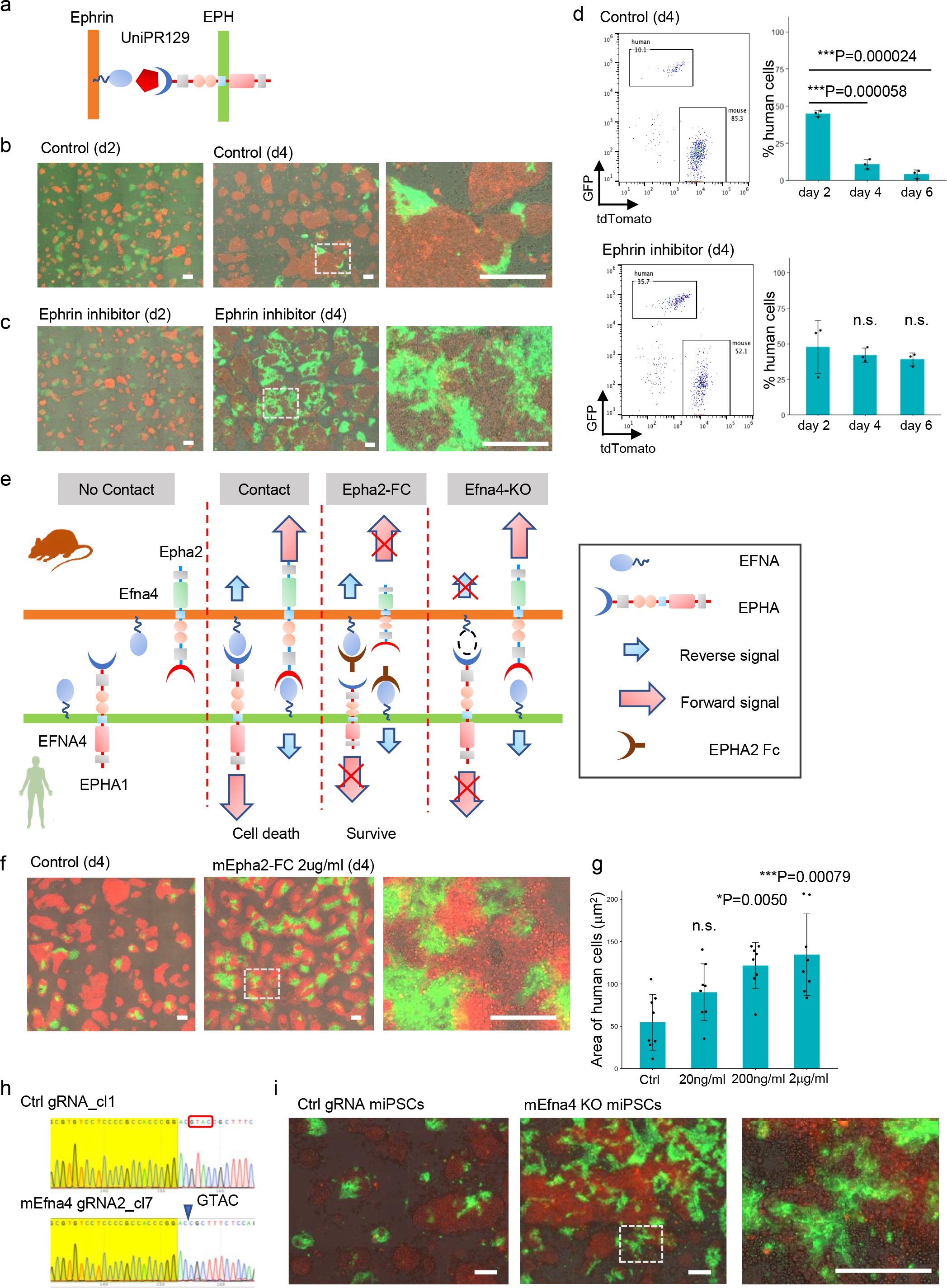
Ephrin forward signaling from mouse cells to human cells regulates interspecies cell competition. **a**. The pan-antagonist of Eph receptors, UniPR129, that binds in Eph receptor, to inhibit whole Ephrin -Eph signaling. **b**, **c**. Representative fluorescence images of day 2 (left) and day 4 (middle and right) co-cultured mGFP-expressing human iPSCs and ntdTomato-expressing moue iPSCs with DMSO (b) or with Ephrin inhibitor (UniPR129) (c). Scale bars, 200Lμm. **d**. Flow cytometry is used to measure the ratio of human cells to human and mouse cells during co-culture with or without the presence of an Ephrin inhibitor. **e**. Conceptual diagram of cell-cell contact-dependent signaling mediated by Ephrin in interspecies cell competition. Forward signaling is inhibited by Epha2-FC, and both forward signaling in human cells and reverse signaling in mouse cells are inhibited by mouse Efna4-KO. **f**. Representative fluorescence images of day 4 co-cultured mGFP-expressing human iPSCs and ntdTomato-expressing moue iPSCs with or without the presence of Epha2-FC protein. The white dashed square indicates a colony where different species of cells are mixed. Scale bars, 200Lμm. **g**. Quantitative results of the changes in the area of human cells on day 4 of co-culture due to varying concentrations of Epha2-FC protein. **h**. Sanger sequencing result showing out-of-frame homozygous 4-bp deletion in Efna4 KO mouse iPSCs. The yellow highlight indicates the PAM region and gRNA region. **i**. Representative fluorescence images of day 4 co-cultured mGFP-expressing human iPSCs and control gRNA moue iPSCs (left) or Efna4 KO mouse iPSCs (middle and right). The white dashed square indicates a colony where different species of cells are mixed. Statistical analysis in d and f: one-way: analysis of variance (ANOVA) with the Tukey post hoc test. \**p* < 0.05, \*\**p* < 0.005, \*\*\**p* < 0.001, no significant change: n.s.. Error bars represent mean ± standard deviation (SD).

Ephrin-Eph signaling activates through the bidirectional mode, and mouse and human iPSCs expressed both Ephrin ligands and receptors (Fig. 3e). Forward signaling is from Ephrin ligands to Eph receptors, while reverse signaling is from Eph receptors to Ephrin ligands^15,16^. It was unclear which forward and reverse signaling is critical for interspecific cell competition because both Ephrin ligands and Eph receptors are expressed in mouse and human iPSCs. To determine which forward or reverse signaling is critical for interspecific cell competition, we analyzed the effects of adding Fc fusion proteins of EphA or Ephrin A, known to act as a forward or reverse signaling decoy, respectively^21^. Among the four types of Fc tested, only EPHA2-Fc significantly overcame cell death in human cells (Fig. 3f, g). The inhibitory effect on hiPSC elimination was in an EPHA2-Fc dose-dependent manner (Fig. 3g), suggesting that EPHA2-Fc acts as a decoy that inhibits the forward signal, potentially both from mouse to human and from human to mouse (Fig. 3e). This indicates that Ephrin forward signaling plays a pivotal role in interspecies cellular competition between the iPSCs. To identify which species’ Ephrin forward signaling is functional, we established Efna4 knockout mouse iPSCs via lentivirus-mediated CRISPR-Cas9 genome-editing in mouse iPSCs (Fig. 3h and Extended Data Fig. 4a-c). In the co-culture of Efna4 KO mouse iPSCs and mGFP-positive human iPSCs, it was shown that by day 4, human cells had a significantly higher survival rate in co-culture with Efna4 KO mouse iPSCs compared to control gRNA- introduced mouse iPSCs (Fig. 3i). These results suggest that in interspecies cellular competition, Efna4 of mouse iPSCs binds to EPHA2 of human iPSCs, leading to the induction of apoptosis, with mouse iPSCs emerging as the winners and human iPSCs as the losers (Fig. 3e). This elucidates the mechanism by which interspecies cell competition is instigated, with mouse iPSCs outcompeting human iPSCs.

### Suppression of Ephrin forward signaling enhances the survival of donor human iPSCs during the chimera animal development process

Ephrin ligands are highly redundant in their expression in various tissues during early mouse development (Extended Data Fig. 5). This highly repetitious expression pattern can most likely mask the single gene knockout phenotype of each ligand and receptor by functional compensation of the ligands and receptors. To avoid this issue and investigate whether inhibition of Ephrin forward signaling can overcome interspecies cell competition in the *in vivo* chimera animal development process, we established mGFP^+^ human iPSCs constitutively expressing EphA2-Fc decoy. For the establishment, we constantly expressed a fusion protein of the EphA2 extracellular domain, with its intrinsic signal sequence replaced by the Il-2 signal sequence, and the human IgG1 Fc region under the CAG promoter (Extended Data Fig. 4d). In co-culture of mGFP^+^ human iPSCs overexpressing EphA2-Fc with tdTomato^+^ mouse iPSCs, EphA2-Fc overexpressing human iPSCs survived significantly better than control mGFP human iPSCs on day 4 (Fig. 4a). Based on this, we examine the effect of Ephrin forward signaling inhibition *in vivo* using human iPSCs expressing EphA2-Fc. We cultured human iPSC under standard primed culture conditions and injected wild-type or EphA2-Fc expressing human iPSCs into wild-type mouse blastocysts (Fig. 4b). At the equivalent of embryonic day 9 (E9), the analysis revealed no GFP signal in mouse embryos injected with control wild-type mGFP human iPSCs (Fig. 4c). However, in one out of six mouse embryos injected with EphA2-Fc human iPSCs, a GFP signal was detected (Fig. 4d). Histological analysis revealed the presence of human mitochondria-positive human cells in the gut and tail region of the mouse fetus, indicating that primed human cells had survived in mouse (Fig. 4e). These results suggest that Ephrin forward signaling, identified in the *in vitro* interspecies primed PSC co-culture system, also controls interspecies cellular competition in the *in vivo* chimera animal development process (Fig. 4f). Since Ephrin-Eph signaling is highly conserved amongst various organisms (Fig. 4g and Extended Data Fig. 6), our study reveals an evolutionarily conserved surveillance system mediated by host Ephrin ligand and human Eph receptor pairs that act as an interspecific barrier during early embryogenesis.

**Figure 4.**
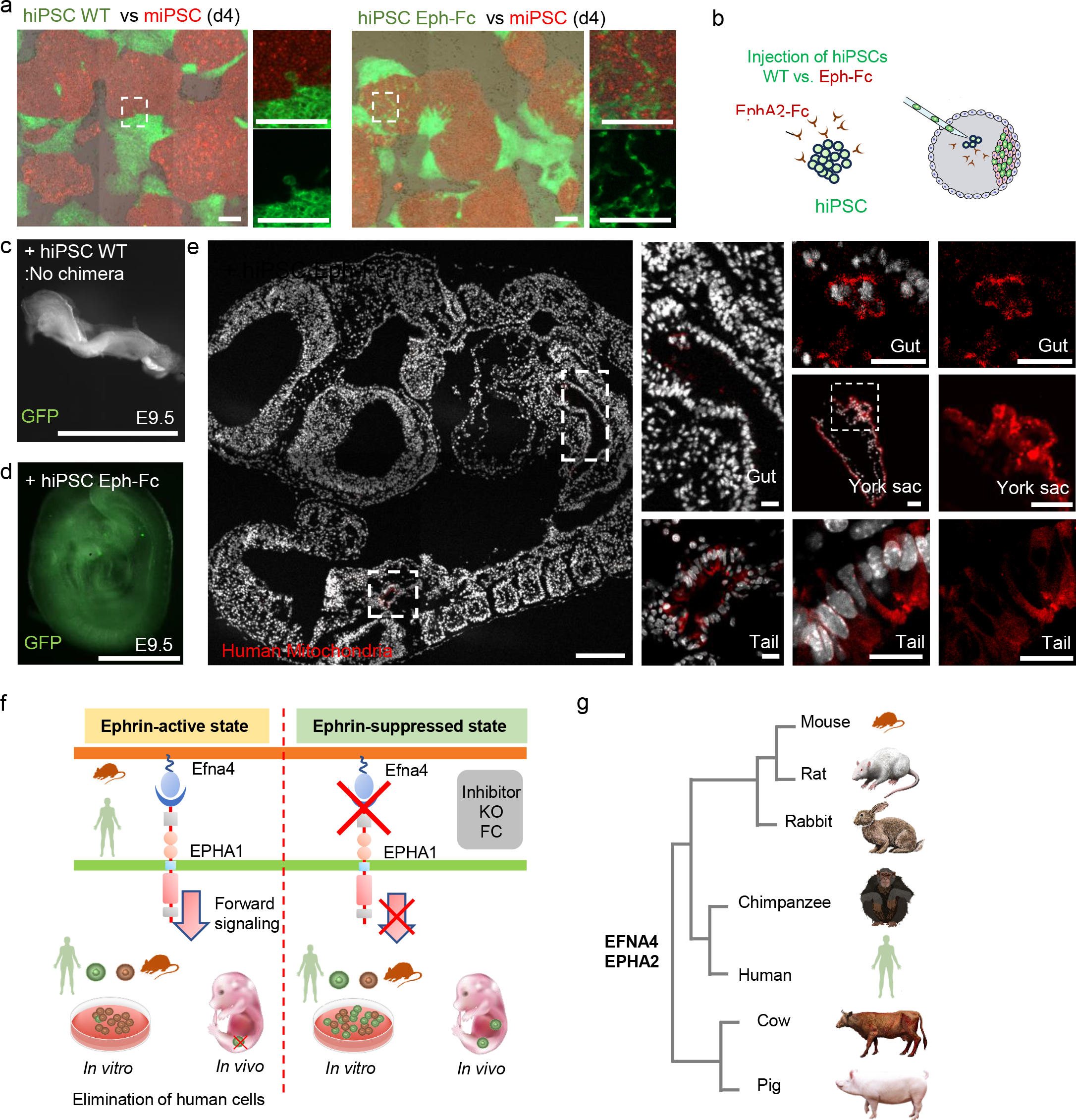
Suppression of Ephrin forward signaling enhances the survival of donor human iPSCs in blastocyst injection. **a**. Representative fluorescence images of day 4 co-cultured ntdTomato-expressing moue iPSCs and WT human PSCs or Epha2-FC expressing human iPSCs. The white dashed square indicates a colony where different species of cells are mixed. Scale bars, 200 μm. **b**. Schema of interspecies blastocyst injection using Epha2-FC expressing human iPSCs. **c**. Representative brightfield and fluorescence merged images of mouse embryos at E9.5 after blastocyst injection with wild-type human iPSCs. Scale bars, 1 mm. **d**. Representative brightfield and fluorescence merged images of mouse embryos at E9.5 after blastocyst injection with Epha2-FC expressing human iPSCs. Scale bars, 1 mm. **e**. Immunofluorescence image of mouse embryos at E9.5 after blastocyst injection with Epha2-FC expressing human iPSCs. Human mitochondria-positive cells were detected in tail, gut, and york sac. Scale bars, 200 μm (left) and 50 μm (right). **f**. Schema of suppression of interspecies cell competition through control of Ephrin signaling. Ephrin forward signaling, identified in the in vitro interspecies primed PSC co-culture system, also controls interspecies cellular competition in the in vivo chimera animal development process. **g**. Phylogeny evolutionary map of EFNA4 and EPHA2.

## Materials and Methods

### Mouse

C57BL/6NCrl mice (cat# 027) as hosts were purchased from Charles River for blastocyst injection. All animal procedures were performed per the ethical guidelines of the Columbia University Medical Center. The animal protocol was reviewed and approved by the Columbia University Institutional Animal Care and Use Committee (IACUC) (protocol: AABF8554). All experiments followed the 2021 Guidelines for Stem Cell Research and Clinical Translation released by the International Society for Stem Cell Research (ISSCR). All human–mouse in vivo interspecies chimeric experimental studies were reviewed and approved by the Columbia University Human Embryonic and Human Pluripotent Stem Cell Research Committee.

### Culture of iPSCs and ESCs

We cultured mouse iPSC (CD1 iPSC or nuclear tdTomato (ntdTomato) iPSC^24^) in 2i/LIF medium on a feeder for maintenance. These mouse PSCs were passaged at a split ratio of 1:10 every 2–3 d. TD1 human iPSCs (Passage#: P15∼P25) in this study were generated from deidentified commercially available human tracheal epithelial cell lines via the manufacturing protocol of Sendai virus-mediated reprogramming (CytoTune2.0) (Thermo Fisher) as described before^24^. Human iPSCs were maintained in feeder-free conditions on laminin iMatrix-511 silk E8 (Amsbio) in StemFit 04 complete medium (Amsbio), supplemented with Primocin (Invivogen), and passaged with TrypLE Select (Gibco) every 1 week. All human iPSC lines screened negative for mycoplasma contamination every other month using a MycoAlert PLUS detection kit (Lonza, LT07-710).

### Mouse iPSC establishment

We established mouse iPSC lines (CD1 iPSC or ntdTomato iPSC) using Sendai virus-mediated reprogramming kit: CytoTune2.0 (Thermo Fisher, A16517) and followed manufactured protocol as described before^24^. Briefly, we harvested E14.5 lung tissues of Rosa26^nt-ng/nt-ng^ mice (Jax, 023035) or CD1 mice (Charles River, 022). The lung fibroblast was maintained in Mouse Embryonic Fibroblast (MEF) medium on a gelatin (Millipore-Sigma, ES006B)-coated well of 6-well plates (0.1 million cells per well). After the three passages of these fibroblasts, we reprogrammed these cells with Sendai virus expressing Yamanaka factors using CytoTune2.0 and cultured them in a 2i/LIF medium^25,26^. Stable iPSC cell line (P7) was established by confirming the negative expression of Sendai virus and positive expression of pluripotency markers (Sox2, Oct4, Nanog, and Ssea1) by immunostainings. To establish ntdTomato^+^ iPSCs, GFP^-^tdTomato^+^ live cells were sorted out by SONY MA900, and single clones were expanded. These colonies were expanded from a single colony, expanded and switched the medium from 2i/LIF to a2i/LIF medium around passage 10, and maintained in a2i/LIF medium afterward, and stocked at P10 for the downstream analyses. Passage number (ntdTomato miPSC: P15∼P20; CD1 miPSC: P15∼P20) 2i/LIF-cultured miPSC were used for this study. All mouse iPSC lines screened negative for mycoplasma contamination every other month using a MycoAlert PLUS detection kit (Lonza, LT07-710).

### Primed PSC culture

Mouse and human primed PSCs were either cultured on MEFs in NBFR medium, as previously reported^5^, which contains DMEM/F12 (Invitrogen) and Neurobasal medium (Invitrogen) mixed at 1:1 ratio, 0.5× N2 supplement (Invitrogen), 0.5× B27 supplement (Invitrogen), 2 mM GlutaMax (Gibco), 1× nonessential amino acids (NEAA, Gibco), 0.1 mM 2-mercaptoethanol (Sigma-Aldrich), 20 ng ml^-1^ FGF2 (PeproTech), 2.5 μM IWR1 (Sigma-Aldrich), and 1 mg mL^-1^ BSA (low fatty acid, MP Biomedicals).

### Surface EGFP construct and lentiviral-mediated gene transduction

The lentiviral vectors pHAGE-EF1a-Cre-w41 were kindly gifted from Dr Darrell N. Kotton, Boston University. To express surface EGFP (sGFP) or membranous EGFP (mEGFP), we amplified IgK signal peptide and PDGFR transmembrane domain by PCR from the template of pcDNA3.1-kappa- myc-dL5-2xG4S-TMst (Addgene, 73206) or pCA-mTmG (Addgene, 60953), respectively. These fragments were cloned into pHAGE-EF1a-Cre-w41 plasmid using an infusion cloning kit (Takara, 638947). Lentiviral vectors were transfected into the packaging HEK293 cell line. After packaging the virus, the supernatant was concentrated by ultracentrifugation as described before^23^. We performed lentiviral-mediated gene transduction to human iPSCs. sEGFP- or mEGFP- stably expressing iPSCs were established after the single colony expansion post-sorting by FACS (SONY MA900) in a complete StemFit medium.

### Generation of i-Switch mouse PSCs

To express a-GFP-NotchECD-TM-TetR-VP64 in mouse CD1 iPSCs, we amplified EF1a promoter region from pEP4 E02S CK2M EN2L by PCR and cloned into pHR_SFFV_LaG17_synNotch_TetRVP64 using infusion cloning kit (Takara, 638947). After packaging the lentivirus, we performed lentiviral-mediated gene transduction to mouse iPSCs. Myc-tag-positive cells were collected by FACS at 72 hours after infection and plated in a2i/LIF medium. Single clones were picked and expanded. To express a-GFP-NotchECD-TM-TetR-VP64 in mouse PSCs, we cloned PGK promoter puromycin resistance region into pLV_tetO_mCherry. After packaging the lentivirus, we performed lentiviral-mediated gene transduction to a-GFP-NotchECD- TM-TetR-VP64 expressing mouse iPSCs. Infected cells were selected by Puromycin for 1 week. Single clones were picked and expanded and maintained in 2i/LIF medium.

### CRISPR knockout

The sequences of sgRNAs are included in Supplementary Table 3. Three sgRNAs were cloned into the LentiCRISPRv2-mCherry plasmid by ligating annealed oligonucleotides with BsmbI-digested vector. The plasmid carrying the specific sgRNA was then transfected into mouse iPSCs using an electroporator (Neon). mCherry-positive cells were collected by FACS at 48 h after transfection and plated. Single clones were manually picked up, expanded. Homozygous knockout clones were confirmed by Sanger sequencing, and the established EphrinA4 knockout CD1 miPSC (P20∼25) was used for this study.

### Generation of Epha2-FC expressing human PSCs

To express Epha2-FC chimeric protein in human PSCs, we amplified IL2 signal peptide from pFUSE-rIgG-Fc2-ICAM1, mouse Epha2 extracellular domain without endogenous signal peptide from mouse genomic DNA, GS linker human IgG1 FC region from pcDNA3-sACE2v2-Fc (IgG1) and PGK promoter puromycin resistance region cloned into lenti CAG-FLAG-dCas9-VPR using infusion cloning kit (Takara, 638947). After packaging the lentivirus, we performed lentiviral-mediated gene transduction to human iPSCs. Infected cells were selected by puromycin for 1 week. Single clones were manually picked and expanded in a complete StemFit medium.

### Interspecies co-culture

For human–mouse co-culture experiments, we used PSCs that had been cultured in a primed culture condition for at least two passages. Human and mouse PSCs were seeded at 4:1 ratio at 2 × 10^4^ cells cm^-2^ on feeder cells in gelatin-coated dish. The cell ratios at each indicated time point were measured by flow cytometry using EGFP expression and Epcam-BV421 (1:50).

### Immunofluorescence

Before the immunostaining, antigen retrieval was performed using Unmasking Solution (Vector Laboratories, H-3300) for 10 min at around 100 °C by microwave. 4-μm tissue sections were incubated with anti-human mitochondria antibodies (1:500) (Abcam; ab92824) overnight at 4 °C. And incubated with secondary antibodies conjugated with Alexa 488 or 567 (ThermoScientific, 1:400) with DAPI for 1.5 h, and mounted with ProLong Gold antifade reagent (Invitrogen, P36930). The images were captured by a Zeiss confocal 710 microscopy.

### Real-time RT-PCR

Total RNA was extracted using a Direct-zol RNA MiniPrep Plus kit (Zymo Research, R2072), and cDNA was synthesized using Primescript RT Master Mix (Takara, RR036B). The cDNAs were then used as templates for quantitative RT-PCR analysis with gene-specific primers. Reactions (10 µl) were performed Luna Universal qPCR Master Mix (New England Biolabs). mRNA abundance for each gene was determined relative to GAPDH mRNA using the 2−ΔΔCt method. Data were represented as mean ± standard deviation of measurements.

### Blastocyst preparation and embryo transfer

Blastocysts were prepared by mating CD1 males with super ovulated CD1 females. Blastocysts were harvested at E3.5 after superovulation. Twenty sorted PSCs were injected into each blastocyst. After the PSC injection, blastocysts were cultured in an M2 medium (Cosmobio) for a few hours in a 37°C, 5% CO2 incubator for recovery. Then, blastocysts were transferred to the uterus of the pseudo pregnant foster mother.

### Cell dissociation for scRNA-Seq

mCherry+, mCherry- iSwitch miPSC, or hiPSC were dissociated with the Tryple Select and stained with DAPI to detect dead cells by the SONY MA900. The live mCherry^+^ or mCherry^-^ Epcam^+^ miPSC or GFP^+^ hiPSC were sorted using the following staining: anti-Epcam antibody (1:50) (Biolegend; 118225). Cells were loaded on a 10x Genomics Chromium controller.

### scRNA-seq data analysis

All scRNA-seq assays were performed with Illumina NovaSeq 6000. Reads alignment to GRCh38 human and GRCm38 mouse reference genome and generation of gene expression matrix were processed using 10x genomics cellranger software version 6.1.2. For processing, integration, and downstream analysis, the Seurat package was used. Cells were selected in the range of 3,000 to 10,000 mapped genes. Feature data were scaled using the Seurat ScaleData function. Non-linear dimension reduction was performed using uniform manifold projection (UMAP) and the Seurat RunUMAP process. The clustering was performed using the Louvain algorithm, and parameters were set empirically by detecting marker genes in each cluster.

### Cross-species scRNA-seq data analysis

For the cross-species cell-cell interaction analyses, we used SCGEATOOL (scTenifoldXct)^18,27^ and CellChat^17^ software. We combined datasets of the mouse (mCherry^+^) and human iPSCs (GFP^+^). By the default parameter of the CellChat analysis, we detected each L-R pair candidate. Since interspecific cell competition occurs exclusively upon cell-cell contact, we selected the L-R pair cell of surface molecules from the mouse or human side and excluded the L-R pair involving secreting proteins. We also analyzed L-R pairs using scTenifoldXct software and ranked the genes in the top 20 hits.

### Statistical analysis

Data analysis was performed using R-studio. Data acquired by performing biological replicas of two or three independent experiments are presented as the mean ± SD. Statistical significance was determined using a student t-test. \**p* < 0.05, ns: non-significant.

## Acknowledgments

We sincerely appreciate the considerate support and scientific input from Dr. Wellington Cardoso at the Columbia Center for Human Development (CCHD) and the members of Cardoso’s lab and CCHD. We acknowledge the support from the CCHD Medicine Microscopy core (MMC), Columbia Stem Cell Initiative (CSCI) Flow Cytometry core (SONY MA900), and Genetically Modified Mouse Model Shared Resource (GMMMSR) for blastocyst injection. This work was funded by NIH-NHLBI 1R01 HL148223-01, DoD PR190557, PR191133 to M. M., JSPS21KK0290, and The Uehara Memorial Foundation to J. T. The work of Jun Wu lab at UT Southwestern Medical Center is supported by the New York Stem Cell Foundation (NYSCF Robertson Investigator Award), Virginia Murchison Linthicum Scholar, and NIH (HD103627- 01A1).

## Extended Data Figure Legends

Extended Data Figure 1. Generation of an interspecies cell-cell contact reporter system.

**a**. Real-time RT-PCR analysis of pluripotent, prime type, and naive genes in the a2iLIF^22,23^ or prime type cultured hiPSCs. The data are normalized to GAPDH and are presented as meansL±Ls.d. (*n*L=L3 biological replicates). Statistical significance was determined by Student-t test. **b**. Representative brightfield and mCherry images of NIH-293T cells infected with previous synNotch system in mono-culture condition. Scale bars, 50Lμm. **c**. Representative mCherry, GFP, and brightfield merged images of the iSwitch system. The mouse iPSCs contacted with sGFP-positive human iPSCs expressed mCherry at 10 hours. Scale bars, 50Lμm. **d**. The ratio of mouse cells and human cells during interspecies co-culture using flow cytometry.

Extended Data Figure 2. Detection of interspecies-specific apoptotic cell populations by scRNAseq.

**a**. Top 20 terms in the GO term enrichment analysis for genes upregulated in cluster 12 compared to all cells. X-axis showing a number of genes. **b**. Feature plots for apoptosis-related genes. Those genes are expressed in cluster 12.

Extended Data Figure 3. Regulation of interspecies cell competition by Ephrin inhibitor.

**a**. Flow cytometry for the ratio of human cells and mouse cells during co-culture with or without the presence of ephrin inhibitor. Human iPSCs were gated by GFP, and mouse iPSCs were gated by tdTomato. **b**. The ratio of mouse cells and human cells during interspecies co-culture using flow cytometry. **c**. Representative brightfield and fluorescence merged images of mouse iPSCs and human iPSCs in monoculture with ephrin inhibitor. Scale bars, 200Lμm.

Extended Data Figure 4. Establishment of Efna4 KO mouse iPSCs by CRISPR Cas9.

**a**. Flow cytometry for lentivirus for CRISPR sgRNA-infected mouse PSCs. Mouse Epcam and mCherry double-positive cells were sorted for single-cell cloning. **b**. The Sanger sequencing result shows out-of-frame homozygous 4-bp deletion (GTAC) in the Efna4 KO mouse iPSC clone. The yellow highlight indicates the PAM region and gRNA region. **c**. Representative bright field and fluorescence images of control or Efna4 KO mouse iPSCs. Efna4 KO mouse iPS cells maintained a morphology similar to the control. Scale bars, 50Lμm. **d**. For Epha2-FC expressing human iPSCs, we generated and infected lentivirus for IL-2 signal peptide-EPHA2 extracellular domain without native signal peptide- human IgG1FC. Infected human iPSCs were selected by puromycin.

Extended Data Figure 5. Ephrin ligand expressions in mouse developmental stages based on deposited scRNAseq database.

**a**. UMAP diagram based on deposited mouse scRNAseq data from E6.5 to E8.5. The annotation results (left) and visualization by developmental time (right) are shown. **b**. Feature plots for Ephrin ligands. Ephrin ligands are highly redundant in their expression in various tissues during gastrulation.

Extended Data Figure 6. Evolutionary phylogeny map of EFNA4 genes.

The EFNA4 gene genomic sequence among species was compared using the Ensembl genome browser. The EFNA4 gene is highly conserved in mammals. Red dotted lines indicate the species used in Figure 4g.

