## Extended Data Figure 1-6 for "Ephrin Forward Signaling Controls Interspecies Cell Competition in Pluripotent Stem Cells"

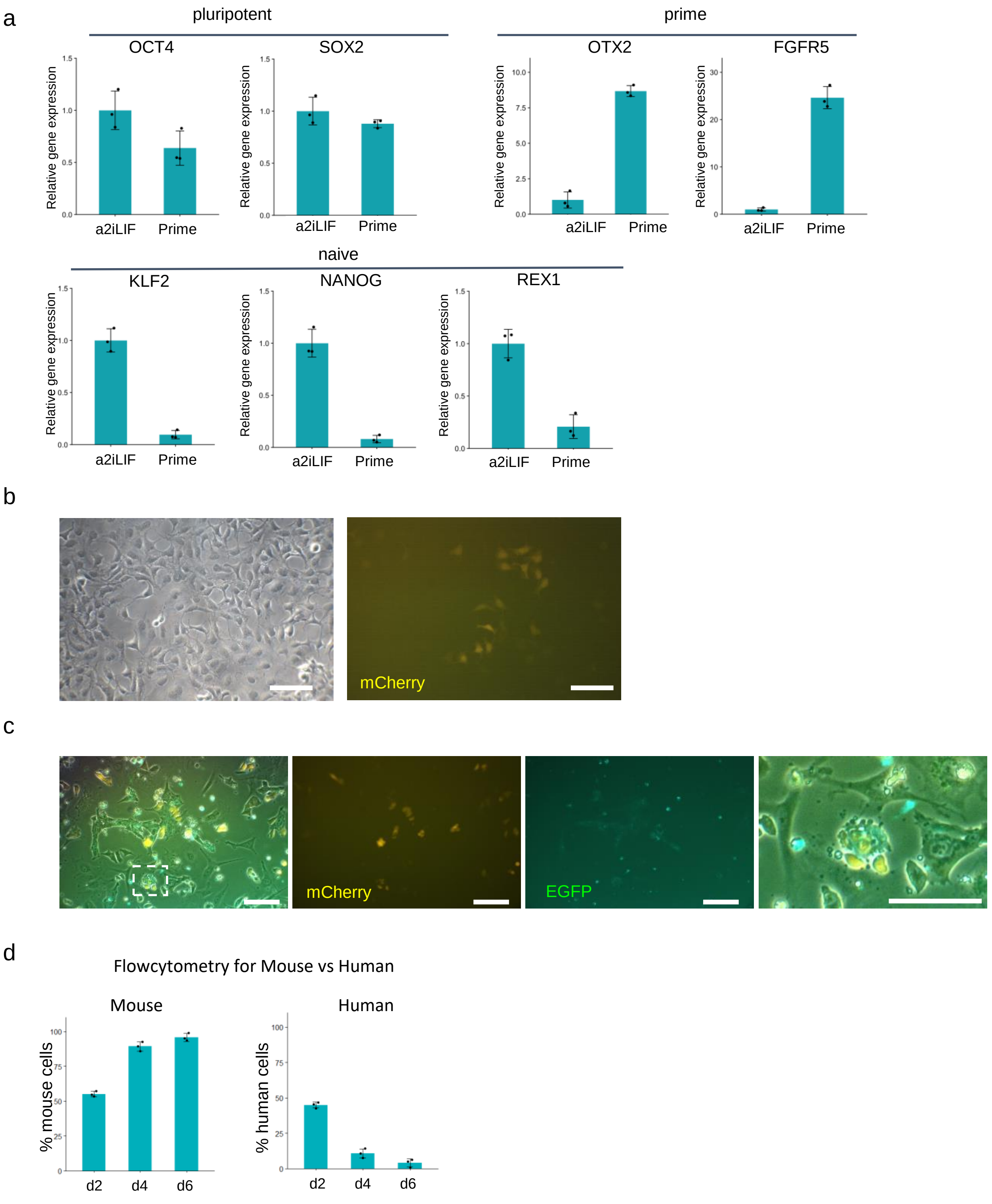


Extended Data Figure 1

a

GO term analysis for C12

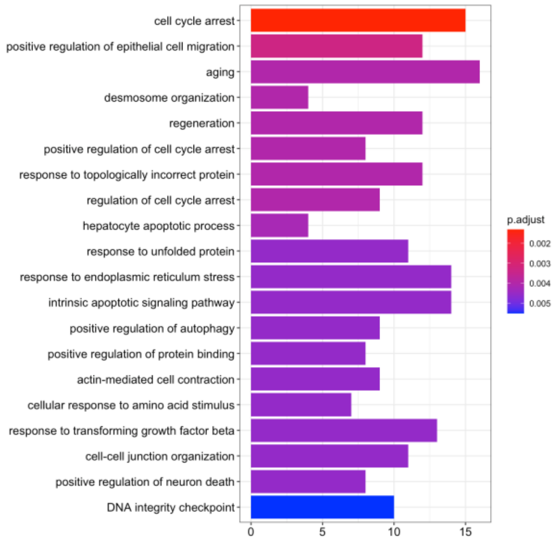


b

Apoptosis related genes expressed in C12

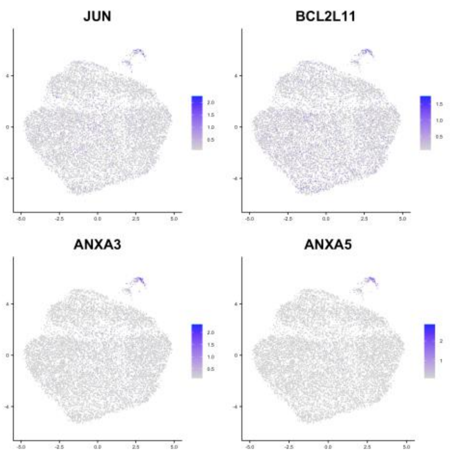


Extended Data Figure 2

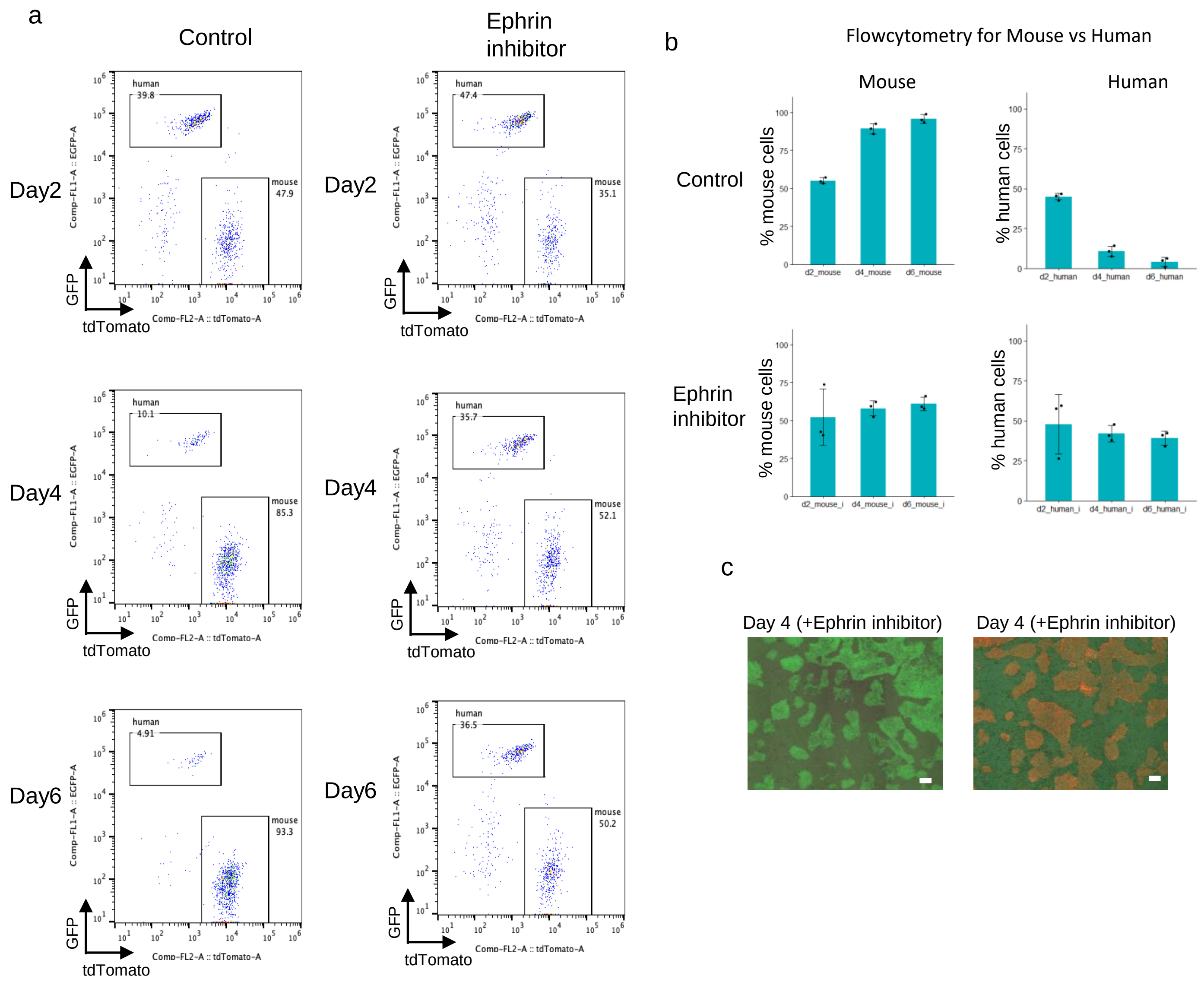


Extended Data Figure 3

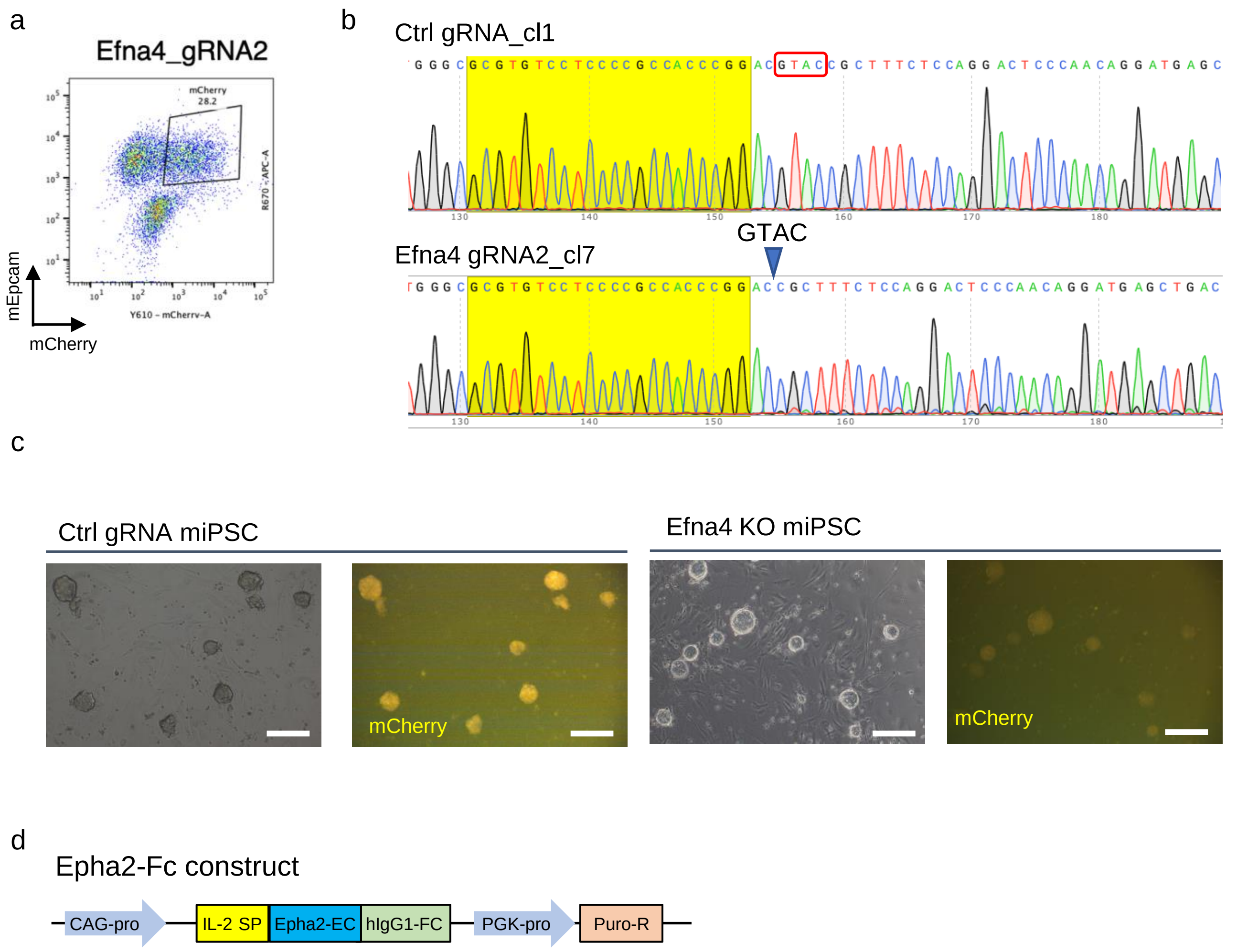


Extended Data Figure 4

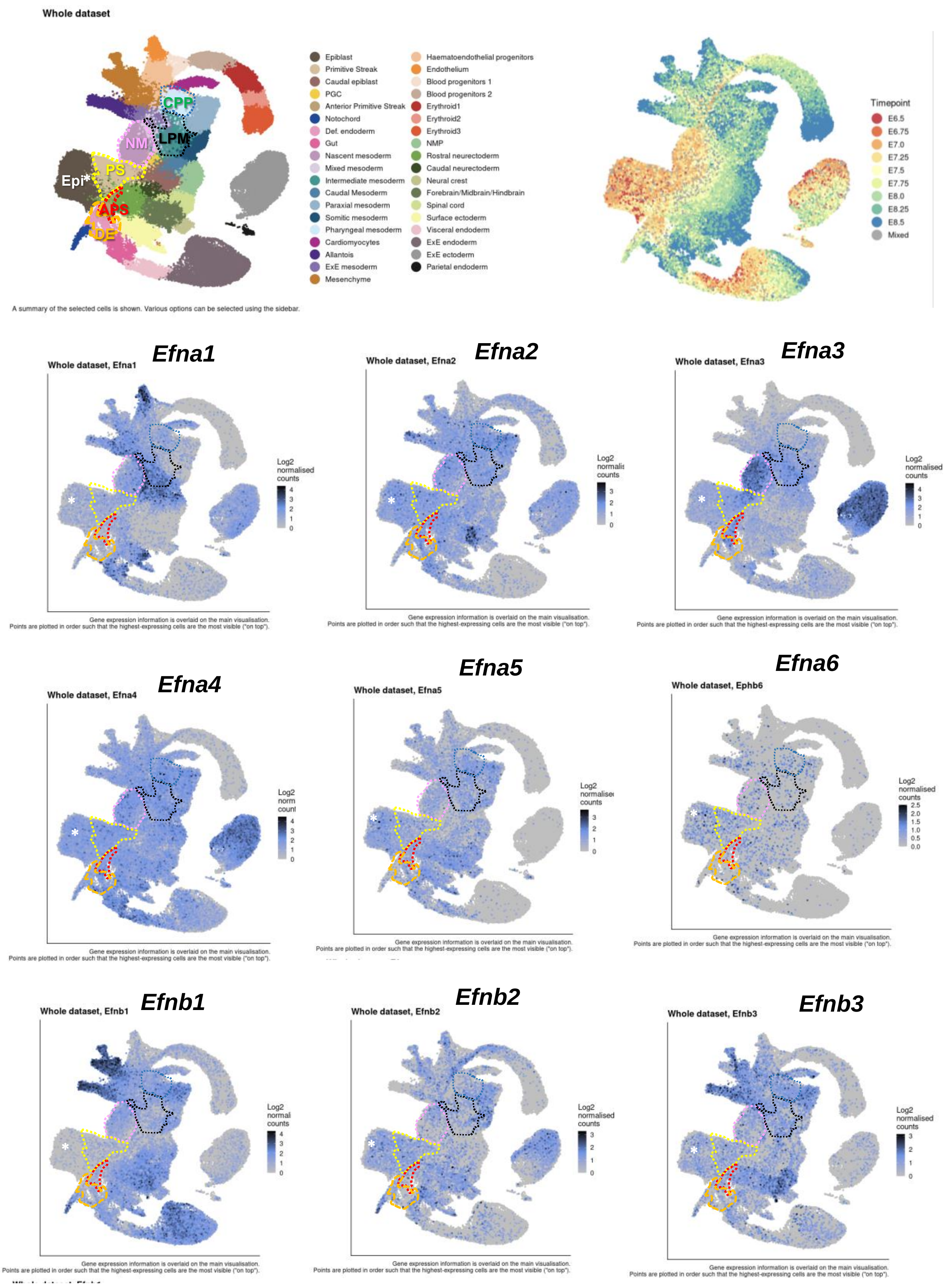


### Extended Data Figure 5

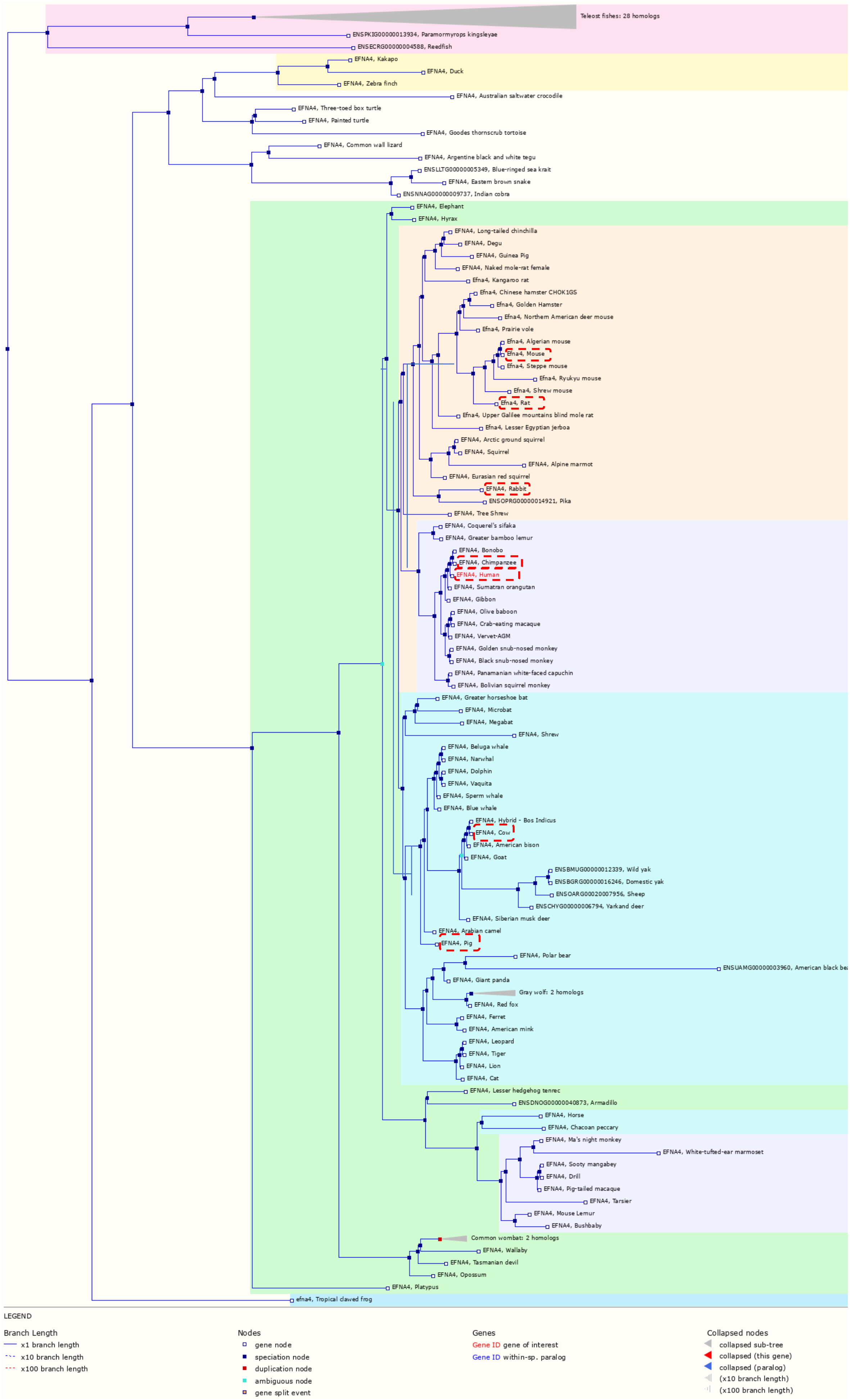


### Extended Data Figure 6
